## Supplemental Figures 1-6 for "Bidirectional redox cycling of phenazine-1-carboxylic acid by *Citrobacter portucalensis* MBL drives increased nitrate reduction"

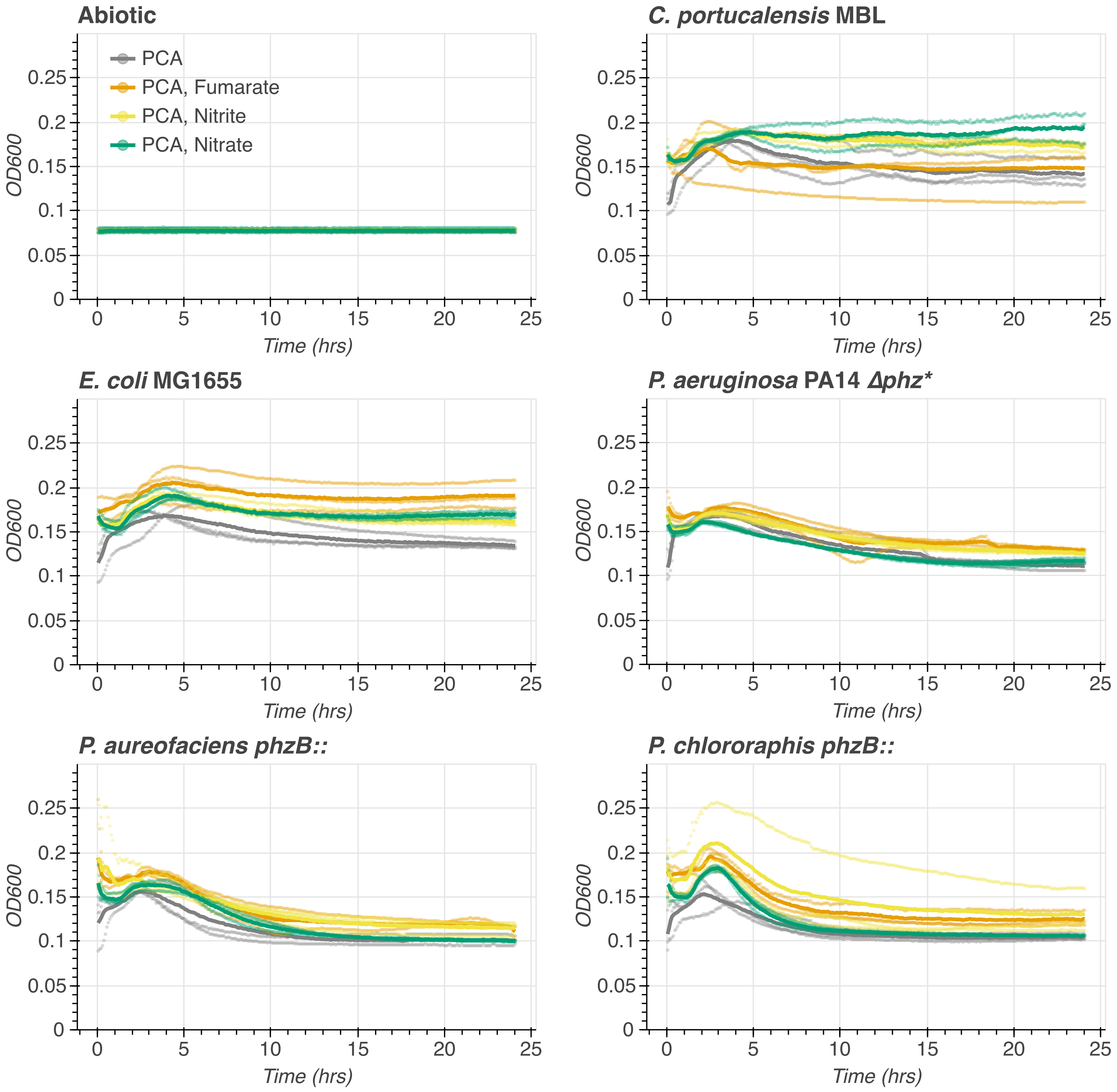


Supplementary Figure 1. OD600 measurements for all strains in all conditions during the oxidation assays (Fig. 1A). Light circles represent all measurements from three independent biological replicates corresponding to the PCAred measurements in Figure 1A. The solid lines are the means of the replicates. The curves are colored by condition (i.e., which, if any, terminal electron acceptor was added). While there are some OD600 trends in the first five hours, there is no evidence of growth that would indicate a doubling. In the abiotic graph, all the data points overlie each other.

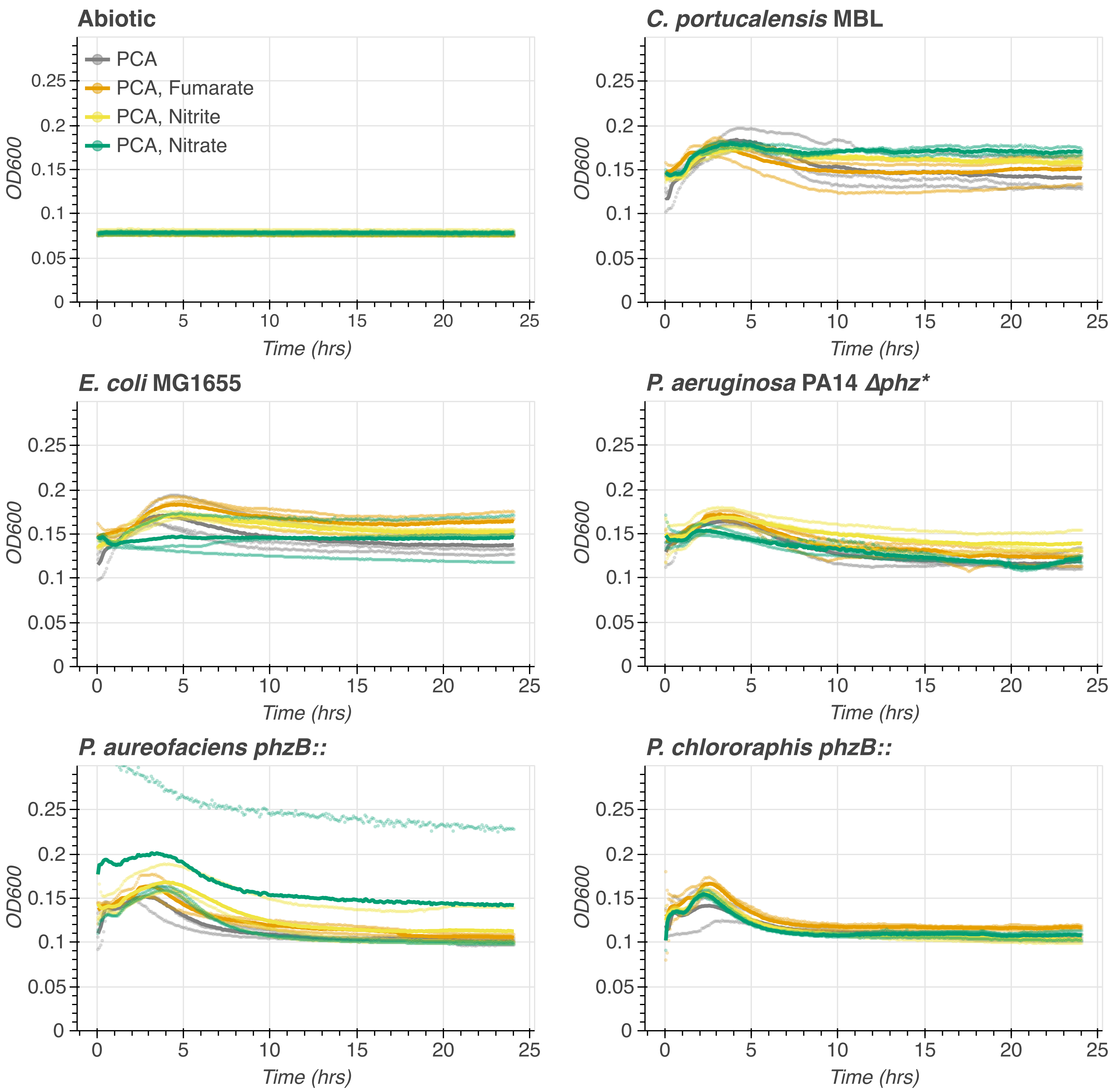


Supplementary Figure 2. OD600 measurements for all strains in all conditions during the reduction assays (Fig. 1B). Light circles represent all measurements from three independent biological replicates corresponding to the PCAred measurements in Figure 1B. The solid lines are the means of the replicates. The curves are colored by condition (i.e., which, if any, terminal electron acceptor was added). While there are some OD600 trends in the first five hours, there is no evidence of growth that would indicate a doubling. In the abiotic graph, all the data points overlie each other.

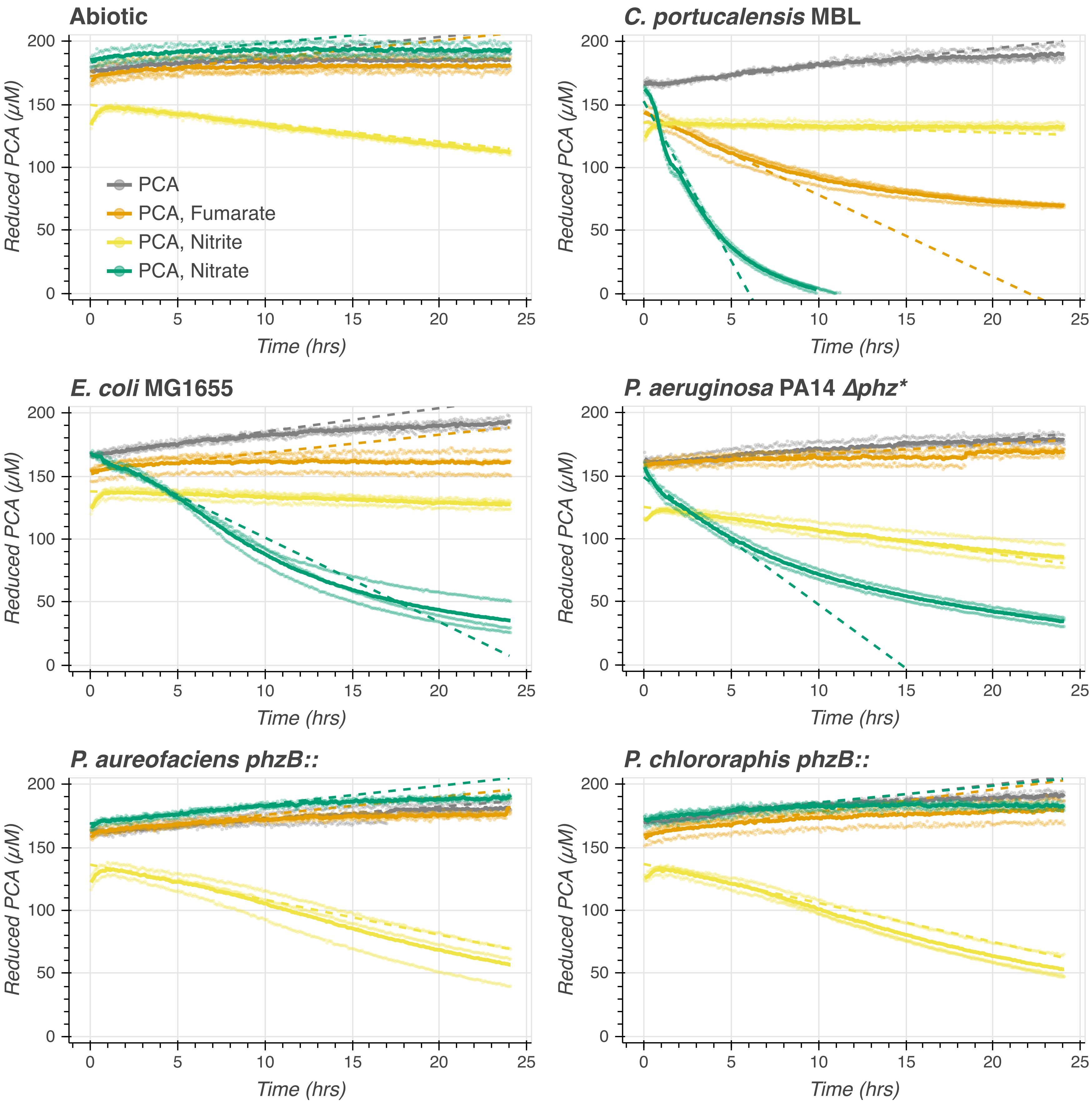


Supplementary Figure 3. Oxidation assay results grouped by strain rather than by condition. This is a different representation of the same data as in Fig. 1A, including the linear regressions (dashed lines) that generated the initial rate estimates as reported in Fig. 1Cα.

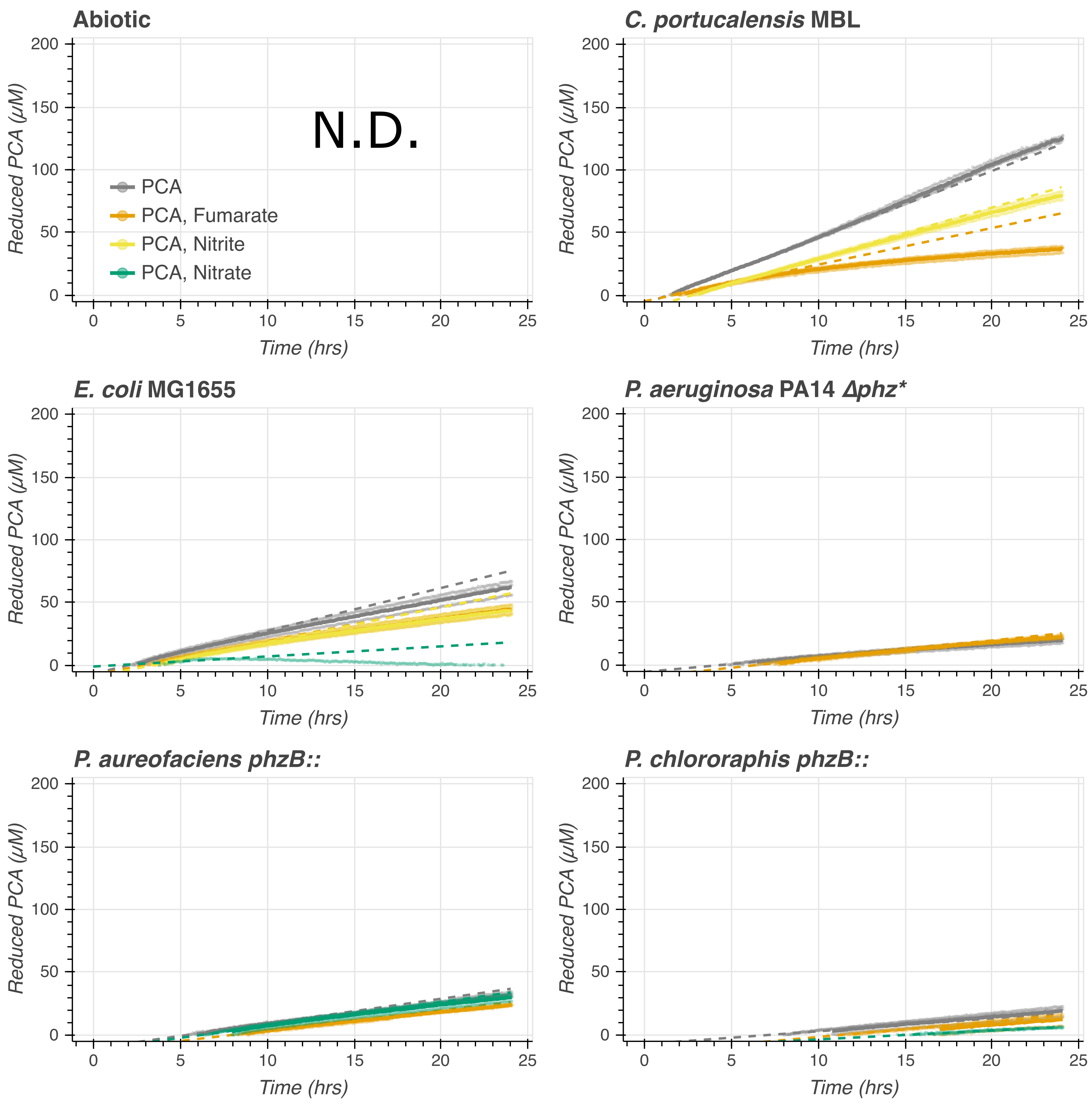


Supplementary Figure 4. Reduction assay results grouped by strain rather than by condition. This is a different representation of the same data as in Fig. 1B, including the linear regressions (dashed lines) that generated the initial rate estimates as reported in Fig. 1Cβ. In the abiotic conditions, no reduction was detected (N.D.).

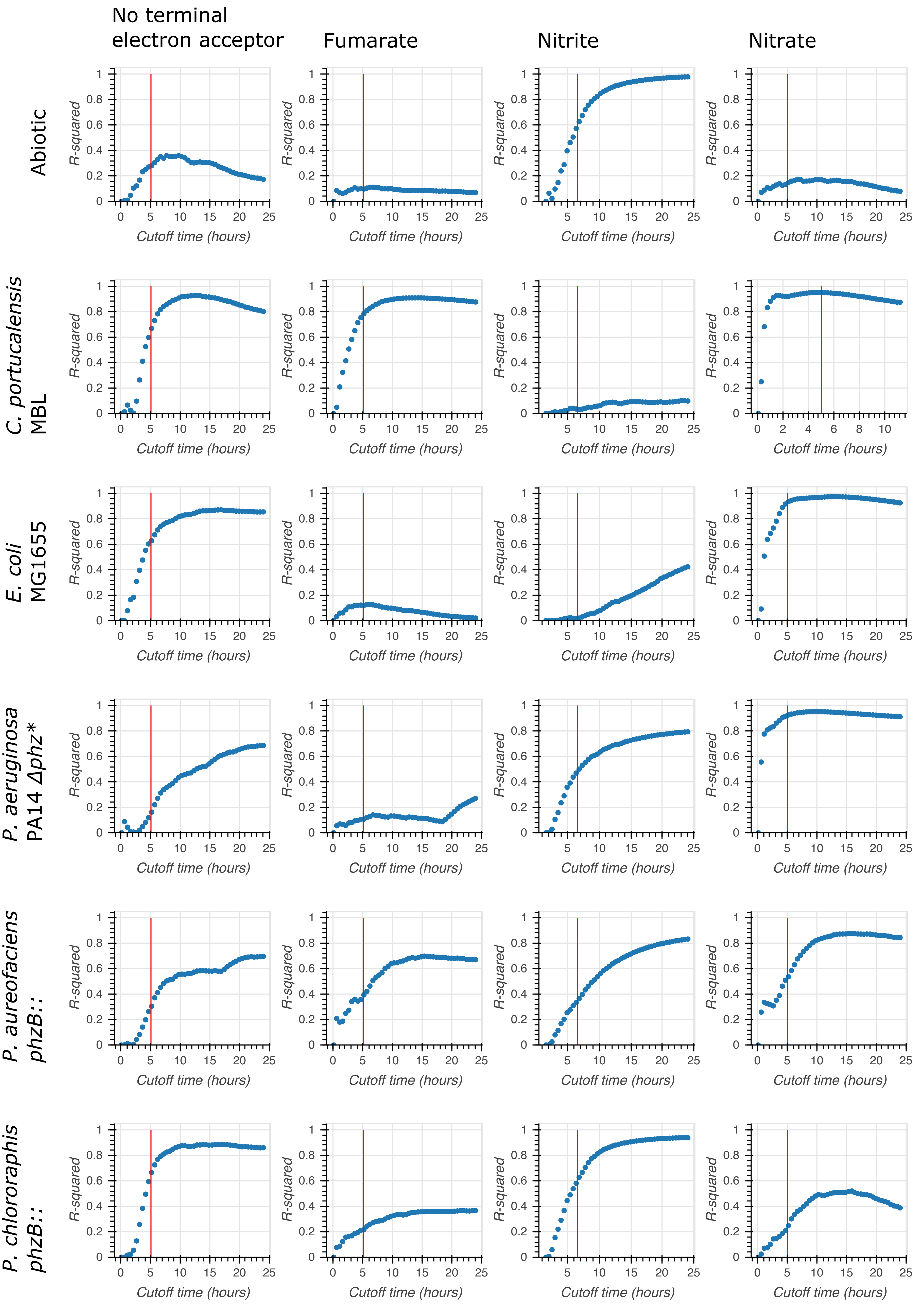


Supplementary Figure 5. Evaluation of the appropriate window for performing the linear regressions to estimate initial redox rates in the oxidation assays (Fig. 1Cα). Starting from the first valid $PCA_{red}$ measurement in each condition, a linear regression was performed over increasing lengths of time and the corresponding $R^2$ value was plotted. A time window of five hours (red line) was selected as a systematic compromise for all conditions.

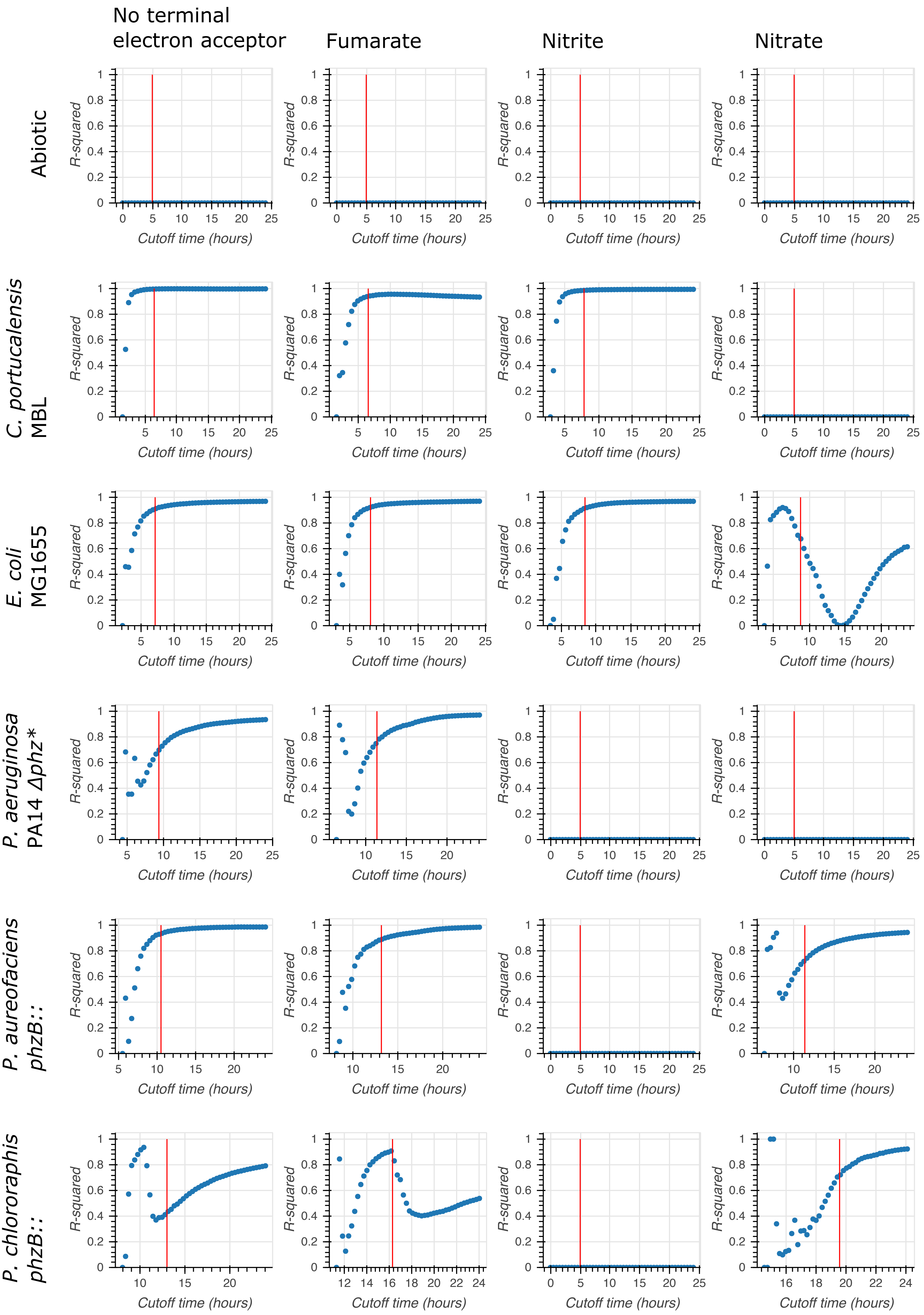


Supplementary Figure 6. Evaluation of the appropriate window for performing the linear regressions to estimate initial redox rates in the reduction assays (Fig. 1Cβ). Starting from the first valid $PCA_{red}$ measurement in each condition, a linear regression was performed over increasing lengths of time and the corresponding $R^2$ value was plotted. A time window of five hours (red line) was selected as a systematic compromise for all conditions.
